# Light-controlled elimination of PD-L1+ cells

**DOI:** 10.1101/2021.08.13.456199

**Authors:** Judith Jing Wen Wong, Pål Kristian Selbo

## Abstract

The programmed death ligand-1 (PD-L1), also known as CD274 or B7-H1, is mainly expressed on cancer cells and/or immunosuppressive cells in the tumor microenvironment (TME) and plays an essential role in tumor progression and immune escape. Immune checkpoint inhibitors (ICIs) of the PD-1/PD-L1 axis have shown impressive clinical success, however, the majority of the patients do not respond to immune checkpoint therapy (ICT). Thus, to overcome ICT resistance there is a high need for potent and novel strategies that simultaneously target both tumor cells and immunosuppressive cells in the TME. In this study, we show that the intracellular light-controlled drug delivery method photochemical internalization (PCI) induce specific and strongly enhanced cytotoxic effects of the PD-L1-targeting immunotoxin, anti-PD-L1-saporin (Anti-PDL1-SAP), in the PD-L1+ triple-negative breast cancer MDA-MB-231 cell line, while no enhanced efficacy was obtained in the PD-L1 negative control cell line MDA-MB-453. Using fluorescence microscopy, we reveal that the anti-PD-L1 antibody binds to PD-L1 on the surface of the MDA-MD-231 cells and overnight accumulates in late endosomes and lysosomes where it co-localizes with the PCI photosensitizer fimaporfin (TPCS_2a_). Moreover, light-controlled endosomal/lysosomal escape of the anti-PD-L1 antibody and fimaporfin into the cytosol was obtained. We also confirm that the breast MDA-MB-468 and the prostate PC-3 and DU-145 cancer cell lines have subpopulations with PD-L1 expression. In addition, we show that interferon-gamma strongly induce PD-L1 expression in the *per se* PD-L1 negative CT26.WT cells and enhance the PD-L1 expression in MC-38 cells, of which both are murine colon cancer cell lines. In conclusion, our work provides an *in vitro* proof-of-concept of PCI-enhanced targeting and eradication of PD-L1 positive immunosuppressive cells. This light-controlled combinatorial strategy has a potential to advance cancer immunotherapy and should be explored in preclinical studies.

## Introduction

Immune checkpoint therapy (ICT) represents a paradigm shift in cancer treatment as it offers durable clinical responses and improves the overall survival of patients with metastatic cancer [1]. ICIs are drugs, e.g., antibodies, blocking the effect of surface inhibitory molecules, including cytotoxic T-lymphocyte-associated protein 4 (CTLA-4) and the PD-1/PD-L1 axis that control the host T cell activity allowing cancers to evade the host immune system [1]. Monoclonal antibodies (mAbs) blocking the PD-1 and PD-L1 interaction can restore effector T cell function, particularly CD8+ T cells, which have a central role in fighting cancer. PD-L1 is a transmembrane protein expressed on the surface of tumor cells and immunosuppressive stromal cells in the TME [2,3]. The PD-L1 expression on tumor cells is not exclusively responsible for the therapeutic effect of PD-1/PD-L1 blockade. Studies have indicated that PD-L1 expression on both malignant cells and immune cells contributes to and modulates T cell dysfunction and immune escape [4]. In addition, Kleinovink et al. demonstrated preclinically that tumors with PD-L1 negative malignant cells could be efficiently treated by PD-L1 blocking antibodies as ICT also targets PD-L1-expressing stromal cells [5]. Despite significant clinical success with PD-1/PD-L1 blockade [6–10], high rates of resistance limit their efficacy, resulting in the majority of cancer patients not responding to this type of cancer immunotherapy [11,12]. It is, therefore, a high need to develop more potent combinatorial targeting strategies that enhance cytotoxic responses in both tumor and immunosuppressive cells.

Burr *et al.* demonstrated that that PD-L1 is taken up into cells by endocytosis [13]. The ability of PD-L1 to undergo internalization, combined with its essential role of suppressing effector T cells, makes it an interesting candidate for targeted delivery of cytotoxic agents [14–16], including type I ribosome-inactivating proteins (RIP). However, a major problem for RIP I protein toxins is entrapment and enzymatic degradation in late endosomes and lysosomes [17,18]. Unlike the highly poisonous type II RIP ricin, saporin lacks the B-chain that facilitates cell surface binding and, in addition, an effective transport mechanism to enter the cell cytosol upon endocytosis. Therefore type I RIPs exhibit low toxicity to intact cells unless the toxin can translocate to the cytosol [28]. It has been hypothesized that RIP-based therapeutics’ cytotoxic effect is caused by a small fraction of the administered dose. The majority of the administered toxin is subjected to lysosomal degradation making RIPs coupled with a targeting moiety attractive candidates to combine with PCI [28–31]. Furthermore, previous studies have indicated that PCI is an efficient method to enhance the delivery of immunotoxins that selectively kill cancer cells expressing CSC markers [30–33]. Interestingly, accumulating evidence suggests an association between PD-L1 expression and cancer stem cell (CSC) phenotype [19–23]. CSCs are a subset of highly aggressive therapy-resistant malignant cells with stem-like features proposed to drive tumor progression, metastasis, and recurrence in patients [24,25]. Moreover, CSCs are suggested to shape the TME into an immunosuppressive landscape [19].

PCI is a modality for cytosolic release of drugs entrapped in endo/lysosomal compartments facilitating enhanced drug efficacy against intracellular targets [26,27]. The PCI technology is based on the use of the amphiphilic photosensitizer TPCS_2a_/fimaporfin, which first resides at the plasma membrane without fully crossing the membrane. Over time, through adsorptive endocytosis, the photosensitizer is accumulated in the membranes of endocytic vesicles that are subsequently transported to endosomes and lysosomes [28]. Upon light exposure at the appropriate wavelength and activation of the photosensitizer leads to reactive oxygen species (ROS) formation. Thus, the PCI method is based on the principles of photodynamic therapy (PDT) [29]. ROS ruptures the membrane of endosomes/lysosomes, thereby releasing the entrapped content that otherwise would have been degraded in the lysosomes [27]. The PCI technology, which was initially developed for soluble macromolecules taken up by endocytosis, has now been evaluated in combination with various therapeutics that accumulate and sequester in endosomes/lysosomes. PCI has been documented as a highly efficient drug delivery method, both *in vitro* and *in vivo*, using therapeutics such as protein toxins, immunotoxins, vaccine antigens (peptides and proteins), and chemotherapeutics such as bleomycin and doxorubicin [26,27]. The clinical relevant PCI photosensitizer, TPCS_2a_/fimaporfin, has been evaluated in a phase I/II combined with bleomycin (Blenoxane®) in cancer patients with advanced and recurrent solid malignancies [30]. Here, PCI of bleomycin was found safe, tolerable, and highly efficient. Currently, TPCS_2a_/fimaporfin-induced PCI of gemcitabine in inoperable bile duct cancer patients is under evaluation in a pivotal phase II trial [31]. Moreover, a phase I study evaluating PCI as a vaccination strategy for peptides (HPV) or proteins (KLH) was recently completed [32]. Hence, preclinical and clinical observations demonstrate that the PCI technology can expand the therapeutic window of drugs and reduce dose-related side effects.

Based on this, we propose that intracellular delivery using PCI of a PD-L1-targeting immunotoxin is an efficient and precision-based strategy to eradicate PD-L1 expressing tumor cells or tumor-associated immunosuppressive cells. The aim of the present study was to evaluate *in vitro* the potential of a PD-L1-targeting immunotoxin combined with PCI as a therapeutic strategy to target and eliminate PD-L1 expressing tumor and immunosuppressive cells. By using flow cytometry and fluorescence microscopy, we show that the triple-negative breast cancer cell line MDA-MB-231 has a high cell surface expression of PD-L1, in line with Latchman *et al.* [33]. Co-localization, observed as yellow fluorescing puncta, of LysoTracker Red and anti-PD-L1-AlexaFluor488 (a green fluorescing mAb substitute for anti-PD-L1-saporin) indicate transport of PD-L1 to late endosomes and lysosomes. Furthermore, the PD-L1-targeting mAb was found to co-localize with the endosome/lysosome-targeting PCI-photosensitizer fimaporfin/TPCS_2a_. After PCI, both anti-PD-L1 mAb and fimaporfin were released into the cytosol. As an *in vitro* proof-of-concept study, we provide evidence that PD-L1 is transported to lysosomes and that PCI of a PD-L1-targeting immunotoxin anti-PD-L1-saporin is specific and efficient in the pico-nanomolar concentrations. Our results suggests a novel light-controlled drug delivery strategy for efficient and safe elimination of PD-L1 expressing immunosuppressive cells in the TME and lays the groundwork for further preclinical explorations.

## Material and methods

### Cell lines and cultivations

The human breast cancer cell lines MDA-MB-231 (ATCC HTB-26), MDA-MB-468 (ATCC HTB-132) and MDA-MB-453 (ATCC HTB-131), the human prostate cancer cell lines PC-3 (ATCC CRL-1435) and DU145 (ATCC HTB-81), the pancreatic cancer cell line PANC-1 (ATCC CRL-1469), and the murine colon cancer cell line CT26.WT (ATCC CRL-2638) were all obtained from the American Type Culture Collection (ATCC, Manassas, VA, USA). The murine colon cancer cell line MC-38 (ENH204-FP) was obtained from Kerafast (Boston, MA, USA). The MDA-MB-231, PC-3, DU145, and CT26.WT cells were maintained in RPMI-1640 culture media (R8758, Sigma-Aldrich, Saint-Louis, MO, USA), whereas MDA-MB-468, MDA-MB-453, PANC-1, and MC-38 were maintained in DMEM (BE12-604F/U1, Lonza Group, Basel, Switzerland). The DMEM media for MC-38 was supplemented with 10 mM HEPES (H0887, Sigma-Aldrich). All culture media were supplemented with 10 % fetal bovine serum (ThermoFisher Scientific, Waltham, MA, USA), 100 U/mL penicillin, and 100 μg/mL streptomycin (Sigma-Aldrich). The cells were mycoplasma negative and maintained in a humidified incubator at 37 °C with 5 % CO_2_.

### Drugs and chemicals

The ICI mAb atezolizumab, a PD-L1 inhibitor (Tecentriq, Roche, Basel, Switzerland), was kindly provided by the Hospital Pharmacy at Oslo University Hospital, Rikshospitalet (Oslo, Norway), and stored at 4 °C. The photosensitizer TPCS_2a_ (fimaporfin) was kindly provided by PCI Biotech AS (Oslo, Norway) and kept in a stock solution of 0.35 mg/mL in polysorbate 80, 2.8 % mannitol, 50 mM Tris, pH 8.5 and stored in dark at 4 °C. MTT (3-(4,5-dimethylthiazol-2-yl)-2,5-diphenyl tetrazolium bromide) (Sigma-Aldrich) for the MTT assay was dissolved in PBS in stock concentration 5 mg/mL and stored at 4 °C in the dark. All work involving TPCS_2a_ was performed under subdued light.

### Immunotoxin

The immunotoxin anti-human PD-L1-saporin (Anti-PD-L1-SAP, BETA-014) was generously provided by Advanced Targeting Systems (ATS, Carlsbad, CA, USA). The targeted toxin is a 295 kDa conjugate between a rabbit polyclonal antibody to human PD-L1 and the secondary conjugate streptavidin-saporin. The non-targeting toxin streptavidin-saporin (IT-27), from here named saporin, was also obtained from ATS.

### Photochemical internalization (PCI) of anti-PD-L1-saporin

MDA-MB-231 (2 x 10^3^ cells/well) and MDA-MB-453 (8 x 10^3^ cells/well) were seeded in 96-well plates (Nunc) and allowed to attach overnight. The cells were then co-incubated with TPCS_2a_ and anti-PD-L1-saporin or saporin as indicated for 18 hours. The TPCS_2a_ concentration was 0.2 μg/mL and 0.4 μg/mL in MDA-MB-231 and MDA-MB-453, respectively. The cells were subsequently washed twice with PBS and chased 4 hours in drug-free medium to remove membrane-bound TPCS_2a_ before illumination. The PCI-photosensitizer fimaporfin/TPCS_2a_ was activated using the LumiSource lamp (PCI Biotech AS, Oslo, Norway) emitting blue light (λ_max_ = 435 nm) as previously described [34].

### Cytotoxicity assay

Viability was evaluated 48 hours post-light exposure using the MTT-assay as previously described [34]. In brief, the cells were incubated with 0.25 mg/mL MTT in culture media for up to 4 hours. The culture media was removed, and the crystals were solubilized using DMSO before the MTT activity (absorbance) was measured at 570 nm [34].

### Flow cytometry

Flow cytometry was performed on live cells to assess the cell surface membrane expression of PD-L1. The cells were detached by trypsin and washed with PBS once. CellTrace™ Violet-stained cells (Thermo Fisher Scientific) were added to all samples prior to antibody staining to serve as an internal control. The samples were blocked with 0.5 % BSA for 10 min on ice and incubated with the primary mAbs biotin-anti-human CD274 (B7-H1, PD-L1) (clone 29E.2A3, BioLegend, San Diego, CA, USA) or biotin anti-mouse CD274 (clone 10F.9G2, BioLegend) in 0.5 % BSA for 20 min on ice. The samples were washed with PBS once and further incubated with streptavidin-AlexaFluor488 (BioLegend) in 0.5 % BSA for 20 min on ice, washed, and immediately analyzed on a flow cytometer LSRII (Becton Dickinson, Franklin Lakes, NJ, USA) as previously described [35]. Data were processed using the FlowJo software version 10.7.1 (Treestar, OR, USA).

### IFN-γ treatment of murine colorectal cancer cells

MC-38 and CT26.WT (1.5 x 10^5^ cells/well) were seeded in 6-well plates (Nunc) and allowed to attach overnight. The cells were then incubated with 100 U/mL mouse recombinant IFN-γ (Invitrogen, Carlsbad, CA, USA) for 24 hours and subjected to flow cytometry analysis of PD-L1 expression as described above.

### Intracellular localization of PD-L1 and TPCS_2a_ before and after light exposure (PCI)

The binding, uptake, and intracellular localization of PD-L1 before and after PCI were evaluated by live-cell fluorescence microscopy in the MDA-MB-231 cells. Cells (3 x 10^5^) were seeded on coverslips in 48-well plates (Nunc), allowed to attach overnight, and washed once with PBS. The cells were then incubated with biotin-anti-human CD274/PD-L1 (BioLegend) on ice for 20 min, washed twice with PBS, and further incubated with streptavidin-AlexaFluor488 (BioLegend) on ice for 20 min. The cells were either evaluated immediately after incubation to visualize the membrane-bound PD-L1 or allowed to internalize for 18 hours. For internalization experiments, fresh culture media containing 0.4 μg/mL TPCS_2a_ was added to the samples and co-incubated with streptavidin-AlexaFluor488. At the end of the incubation (18 hours), the cells were then washed twice with PBS, and chased for 4 hours in drug-free media to remove membrane-bound TPCS_2a_. Image acquisition was performed either pre- or ~1 hour post-blue light exposure (100 seconds). The cells were incubated with LysoTracker™ Red DND-99 (Thermo Fisher Scientific) for 30 min before image acquisition to visualize the acidic organelles. Hoechst 33342 (Thermo Fisher Scientific) was incubated for 15-20 minutes to visualize the nuclei. Image acquisition by a Zeiss Axioplan epifluorescence and phase contrast microscope using the 63x/NA1.4 PlanApo objective (Carl Zeiss AG, Oberkochen, Germany) and image processing and analysis were performed by the AxioVision Analysis software program (Carl Zeiss AG) as previously described [34].

### Statistical analyses

Statistical analyses were performed using two-sided t-test SigmaPlot 14.0 (Systat Software, Inc., San Jose, CA, USA). Statistical significance was set at p ≤ 0.05.

## Results

### Evaluation of cell membrane PD-L1 expression in breast, prostate, and pancreatic carcinoma cell lines

Plasma membrane expression and internalization of PD-L1 is a prerequisite for PCI-based targeting of this protein. First, we therefore screened the PD-L1 surface expression in different human cancer cells; three triple-negative breast cancer cell lines (MDA-MB-231, MDA-MB-468, MDA-MB-453), two prostate cancer cell lines (DU145, PC-3), and one pancreatic cancer cell line (PANC-1) (**Fig. 1A**). To prevent endocytosis, live cells were incubated with the primary and secondary antibody on ice and subsequently subjected for flow cytometry analysis. Several publications indicate high PD-L1 expression in MDA-MB-231 [33,36]. Indeed, our flow cytometry data show a ~ 10-fold increase in median fluorescence intensity (MFI) in the MDA-MB-231 cells compared to the secondary antibody alone (**Fig. 1B**). The minor overlap with the secondary antibody indicates that near 100 % of the cells are PD-L1 positive. A ~ 2-fold increase was detected in both of the prostate cancer cell lines (PC-3 and DU145). In the MDA-MB-453, MDA-MB-468, and PANC-1, the fluorescence intensity was comparable to the secondary antibody alone, indicating low (MDA-MB-468) or very low/no (MDA-MB-453 and PANC-1) PD-L1 membrane expression. As the MDA-MB-231 cells had the highest PD-L1 expression, this cell line was selected for further analysis to provide proof-of-concept of the PCI-based PD-L1-targeting strategy.

**Figure 1.**
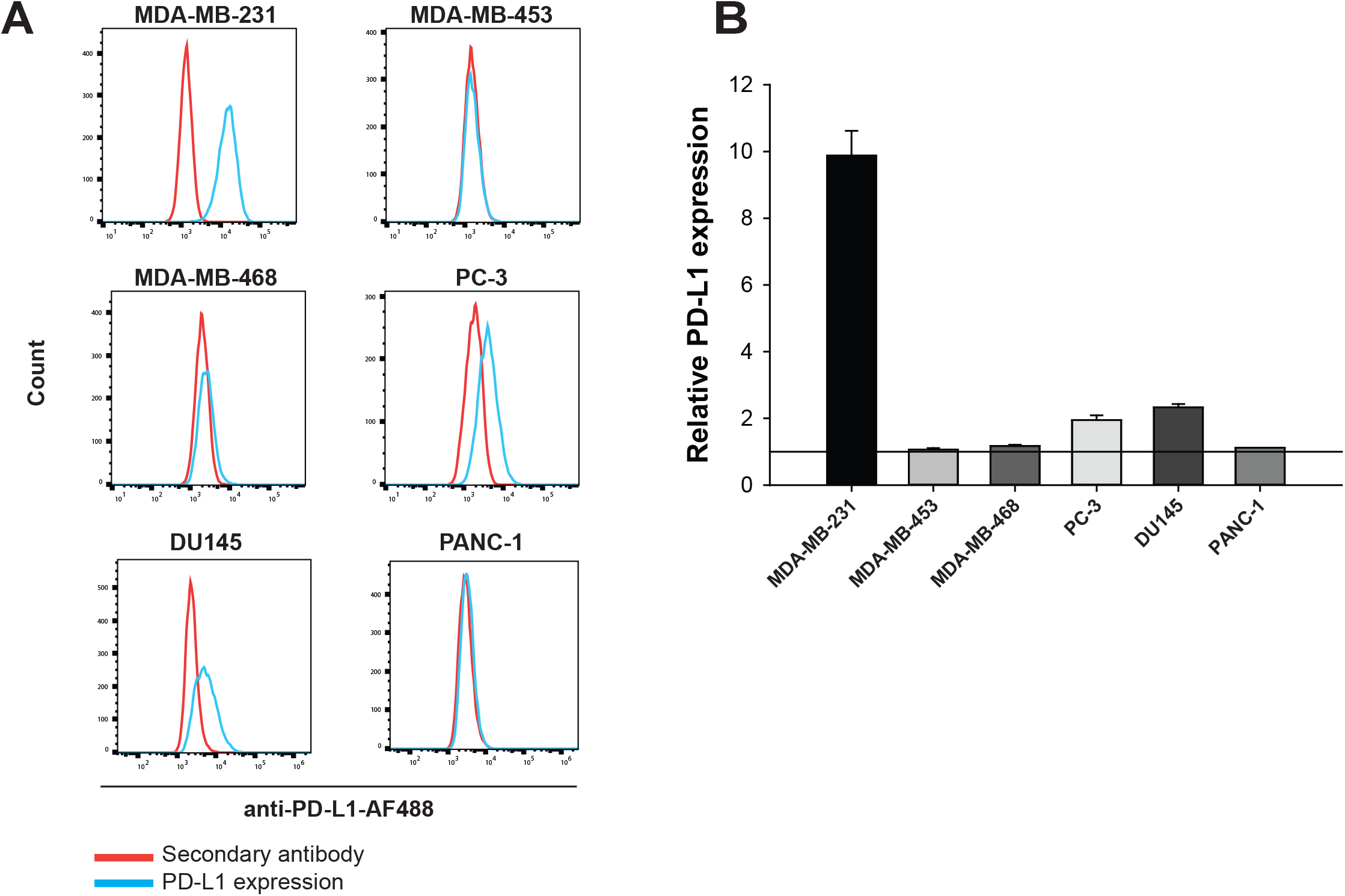
PD-L1 surface expression in the human cancer cell line panel. **(A)** Representative flow cytometry histograms of PD-L1 surface expression on MDA-MB-231, MDA-MB-453, MDA-MB-468, PC-3, DU145, and PANC-1 cells. Red line indicates secondary antibody alone and blue line indicates primary and secondary antibody (PD-L1 expression). **(B)** Quantification of the PD-L1 expression in the cell panel. The relative PD-L1 expression is the median fluorescence intensity of anti-PD-L1-stained sample relative to secondary antibody alone. Data presented as mean ± S.E.M. of at least two independent experiments.

### Anti-PD-L1 mAb binds to PD-L1 on the surface and accumulates in late endosomes and lysosomes

To perform PCI-based intracellular delivery of the anti-PD-L1-targeting immunotoxin, it is crucial that the PD-L1-targeting mAb first binds to PD-L1 and subsequently is taken up by the targeted cells, and localizes to endosomes and/or lysosomes. Thus, surface binding, cellular uptake, and intracellular localization of the PD-L1-targeting mAb (anti-PD-L1-AF488) and its co-localization with LysoTracker Red was first studied using fluorescence microscopy. A similar staining procedure as for flow cytometry was used, and the images were acquired immediately at the end of incubation. Live-cell fluorescence microscopy confirmed the binding of the PD-L1-targeting mAb and thereby PD-L1 expression on the surface of MDA-MB-231 cells (**Fig. 2A**). Image acquisition of the MDA-MB-231 cells 24 hours later revealed fluorescent puncta of anti-PD-L1-AF488 (green), which co-localized (yellow/orange) with the fluorescent marker LysoTracker Red that stains acidic organelles (endosomes and lysosomes) (**Fig. 2B**). Eng *et al.* have previously demonstrated that the photosensitizer fimaporfin/TPCS_2a_ co-localizes with LysoTracker Green in MDA-MB-231 cells [37]. The anti-PD-L1-AF488/PD-L1 is internalized and co-localized with LysoTracker Red in late acidic endosomes and lysosomes over time, which is optimal for performing PCI-based targeting of this immune-checkpoint protein.

**Figure 2.**
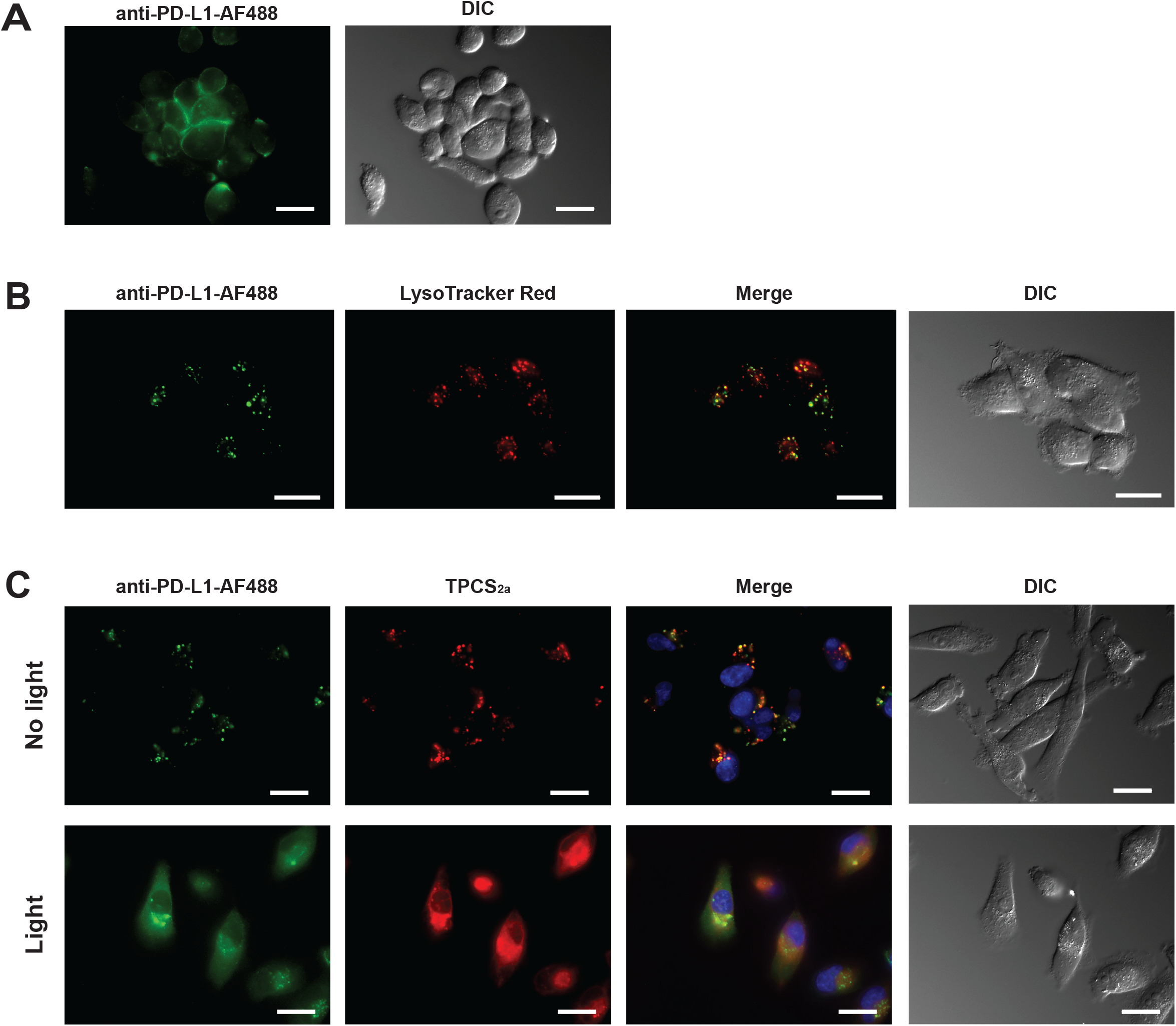
Specific binding, uptake and PCI-induced cytosolic release of PD-L1 antibody. **(A)** Fluorescence microscopy of cell surface binding of anti-PD-L1 antibody (PD-L1-targeting primary antibody bound to secondary antibody labelled with AlexaFluor488, Anti-PD-L1-AF488, green signal) in live MDA-MB-231 cells. The cells were incubated on ice to prevent endocytosis. **(B)** Cellular uptake of anti-PD-L1-AF488 (green) in live MDA-MB-231 cells after overnight incubation (18 hours) at 37 °C and co-localization with LysoTracker Red (red) in live MDA-MB-231 cells. Co-localization of antibody and LysoTracker is indicated in merge (yellow/orange), with corresponding DIC image. **(C)** Upper four panels (No light = before PCI), Anti-PD-L1-AF488 (green) and TPCS_2a_ (red) after 18 hours co-incubation and 4 hours chase,, with corresponding merge and DIC images. Lower four panels (Light = PCI), fluorescence images of PCI-induced cytosolic release of internalized PD-L1 and TPCS_2a_ in live MDA-MB-231 cells. Co-localization (merge) of TPCS_2a_ and anti-PD-L1-AF488 indicated in yellow/orange, with corresponding DIC image. Nucleus (DNA) was stained with Hoechst 33342 (blue). Scale bar = 20 μm. Light exposure (LumiSource): 100 seconds ≈ 0.7 J/cm^2^. All images are representative images of three independent experiments.

### Light-activation and PCI-induced cytosolic delivery of internalized anti-PD-L1 mAb

To achieve endo/lysosomal escape using PCI, which includes photochemical-induced damage of endo/lysosomal membranes, co-localization of the PCI-photosensitizer and the immunotoxin in these vesicles is required in advance of light activation. For that reason, the co-localization of fimaporfin/TPCS_2a_ and anti-PD-L1-AF488 (in these experiments functioning as a substitute for the immunotoxin) in MDA-MB-231 cells before and after PCI was studied using fluorescence microscopy. Overnight incubation of fimaporfin and the PD-L1 antibody, including wash and chase as in the PCI procedure, revealed fluorescent puncta of both drugs with a high degree of intracellular co-localization (yellow/orange signal of merged images) between the anti-PD-L1-AF488 (green) and fimaporfin/TPCS_2a_ (red), indicating co-localization in late endosomes and lysosomes. (**Fig. 2C**). Strikingly, ~ 1 hour post-light exposure, the microscopy images revealed a diffuse fluorescence throughout the cytosol of both fluorochromes, indicating a PCI-induced endosomal/lysosomal escape of both the PD-L1 antibody and the PCI-photosensitizer (**Fig. 2C**).

### Light-enhanced cytotoxic responses of anti-PD-L1-saporin in MDA-MB-231

To demonstrate *in vitro* proof-of-concept, the PD-L1 targeting immunotoxin, anti-PD-L1-saporin, was evaluated in combination with PCI in the PD-L1^high^ expressing MDA-MB-231 cells. Two different PCI protocols were explored; (I) an 18 hours pre-incubation with fimaporfin/TPCS_2a_ prior to a 4 hours incubation of the immunotoxin during the chase period before light exposure. This protocol was compared with the second protocol (II), which included an 18 hours co-incubation of the immunotoxin with fimaporfin/TPCS_2a_ (**Fig. 3A**). Both protocols have previously been evaluated and found to be efficient at enhancing cytotoxic effects of, e.g., CD133 or EGFR-targeting toxins [38,39]. The “4 hours incubation” protocol (I) did not result in any PCI effects, indicating that the cellular uptake of PD-L1 is relatively slow as compared to CD133 and EGFR [38,39]. Strikingly, a light-dose dependent and strongly enhanced cytotoxicity by PCI of 10 pM immunotoxin was observed with 18 hours incubation compared to PCI of the non-targeted control (saporin) (**Fig. 3A**). Using a 10-fold higher concentration of the immunotoxin (100 pM) did not enhance the response in the PD-L1 negative MDA-MB-453 cells compared to PCI of saporin or PDT (**Fig. 3B**).

**Figure 3.**
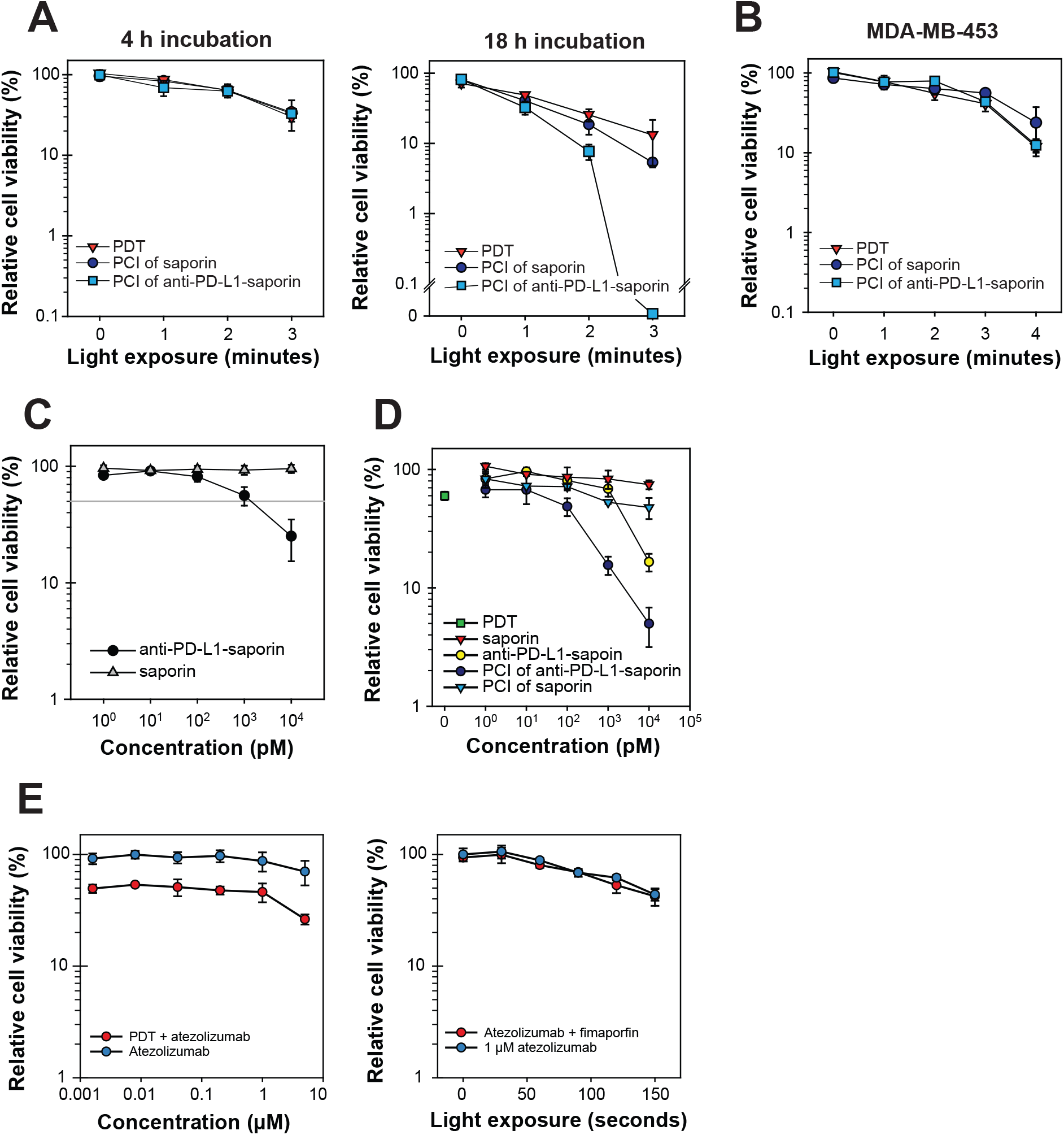
Specific and PCI-enhanced cytotoxicity of anti-PD-L1-saporin in PD-L1^high^ MDA-MB-231 cells. **(A)** Light dose-dependent cytotoxic responses to PCI. Relative cell viability of MDA-MB-231 after PCI of immunotoxin (anti-PD-L1-saporin) or the non-targeting toxin (saporin) compared to PDT (0.2 μg/mL TPCS_2a_ + light). Cells were incubated with 10 pM of the immunotoxin or toxin for 4 (left panel) or 18 hours (right panel) prior to light exposure (1 min = 0.42 J/cm^2^). **(B)** Relative cell viability of PD-L1-negative MDA-MB-453 after PCI of 100 pM anti-PD-L1-saporin or saporin compared to PDT (0.4 μg/mL TPCS_2a_ + light). Representative experiments out of at least two independent experiments. Data presented as mean of triplicates ± S.D. **(C)** Efficacy of immunotoxin and toxin in the absence of PCI. Cytotoxicity of anti-PD-L1-saporin or saporin in MDA-MB-231 cell incubated for 18 hours. The reference line indicates 50 % relative cell viability. Data presented as mean of at least three independent experiments ± S.E.M. **(D)** Concentration-dependent PCI of immunotoxin or toxin in MDA-MB-231 cells using a fixed light dose corresponding to 110 seconds ≈ 0.77 J/cm^2^. **(E)** Atezolizumab (Tecentriq) does not enhance the PDT-efficacy. No enhanced efficacy was achieved by combining PD-L1 blockade with PDT (0.2 μg/mL TPCS_2a_ + light). Left panel, Increasing atezolizumab concentrations in combination with fimaporfin/TPCS_2a_ exposed to 110 seconds light. Right panel, increasing light doses of 1 μM atezolizumab combined with fimaporfin, both co-incubated for 18 hours. For all experiments, cytotoxicity was evaluated 48 hours post-light exposure using the MTT assay. Representative experiments out of at least three independent experiments. Data presented as mean of triplicates ± S.D.

Independent of PCI, the specificity and cytotoxic potential of the immunotoxin, anti-PD-L1-saporin, was further evaluated and compared with the non-targeting toxin saporin. Increasing concentrations of anti-PD-L1-saporin or saporin were incubated for 18 hours in the MDA-MB-231 cells (**Fig. 3C**). The non-targeting toxin did not induce any substantial cytotoxicity (< 10 % viability decrease) over the entire concentration range of 1 – 10,000 pM in the MDA-MB-231 cells. On the other hand, the PD-L1-targeting immunotoxin induced a 44 % and a 75 % reduction in cell viability at 1000 pM and 10,000 pM, respectively. To further validated the specificity and efficacy of anti-PD-L1-saporin, concentration-dependent PCI experiments in MDA-MB-231 cells using a fixed light dose (110 seconds ≈ 0.77 J/cm^2^) corresponding to ~ 50 % reduction in viability with photochemical treatment alone were performed (**Fig. 3D**). Enhanced cytotoxicity was observed after PCI of the immunotoxin. However, compared to photochemical treatment alone, no increased cytotoxic effects was found after PCI of the non-targeting toxin (saporin) across all concentrations, indicating a selective cytotoxic efficacy can be achieved by PD-L1 targeting.

### Blocking PD-L1 with atezolizumab does not enhance PDT efficacy

To evaluate if PD-L1 blockade in MDA-MB-231 cells affected viability or if combining PD-L1 blockade with PDT enhanced the cytotoxicity, we used the clinically approved anti-PD-L1 mAb atezolizumab (Tecentriq®). MDA-MB-231 cells were either incubated with atezolizumab alone or co-incubated with the photosensitizer fimaporfin/TPCS_2a_, subsequently washed, chased, and illuminated as in the PCI protocol. Both increasing atezolizumab concentrations combined with a fixed light dose (110 seconds light exposure ≈ 0.77 J/cm^2^) and increasing light exposure times combined with a fixed atezolizumab concentration (1 μM) were evaluated (**Fig. 3E**). At the highest atezolizumab concentration (5 μM), the viability was reduced by ~ 35 %, whereas lower concentrations (e.g., 100 nM) did not reduce the cell viability. Passariello *et al.* have previously reported a ~ 25 % reduction in viability in MDA-MB-231 cells after 100 nM atezolizumab incubation for 72 hours [40]. In our experimental set-up, we used a considerably shorter (18 hours) incubation time, corresponding to the same incubation time in the PCI protocol. No enhanced fimaporfin efficacy was achieved by combining PD-L1 blockade, indicating that the PCI-enhanced toxicity of anti-PD-L1-saporin is due to the RIP activity of the toxin and not induced by binding of **atezolizumab** to PD-L1.

### IFN-γ induced PD-L1 expression in murine colon cancer cell lines

Freeman *et al.* first demonstrated that IFN-γ induces expression of PD-L1 [41]. The immunotoxin evaluated here is based on a polyclonal antibody recognizing the human PD-L1. To further evaluate the concept in an immunocompetent model, an immunotoxin targeting the murine PD-L1 is necessary. The murine colon cancer cell lines MC-38 and CT26.WT were selected and evaluated for PD-L1 expression using flow cytometry in live cells (**Fig. 4A**). A ~ 2.2-fold increase in median fluorescence intensity indicating PD-L1 expression was detected in MC-38 cells, whereas the PD-L1 expression was very low in CT26.WT cells (1.2-fold) (**Fig. 4B**). An incubation with IFN-γ for 24 hours induced a ~ 2.4-fold (p < 0.001) and ~ 3.4-fold (p < 0.05) higher PD-L1 expression in in the CT26.WT and MC-38 cells, respectively. These results concur with other studies [5,41,42]. Thus, both murine cell lines can be used as a model for a PD-L1 targeting immunotoxin in future studies.

**Figure 4.**
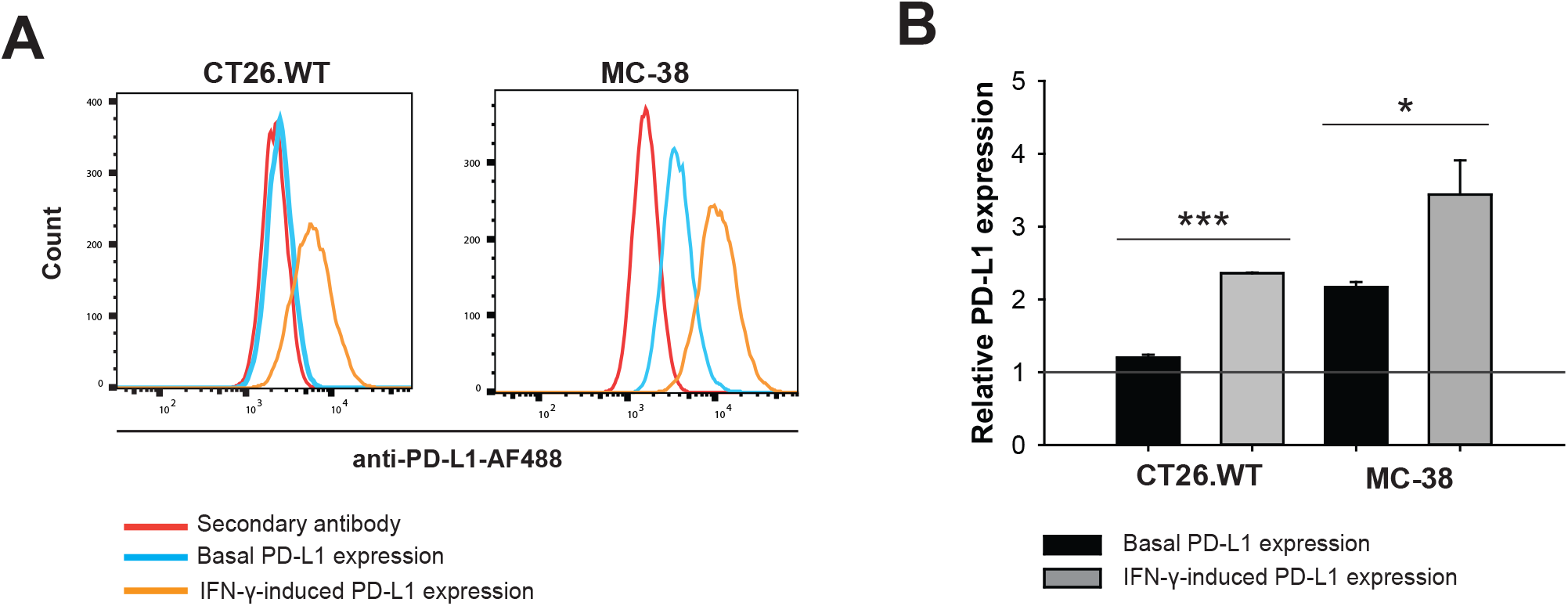
IFN-γ induce surface expression of PD-L1 in murine colon cancer cells. **(A)** Representative flow cytometry histogram of surface PD-L1 expression in the murine colon cancer cell lines CT26.WT and MC-38. Red line indicates secondary antibody alone, blue line indicates primary and secondary antibody (PD-L1 expression), and orange line indicates IFN-γ treated cells (100 U/ml, 24 hours). **(B)** Quantification of the PD-L1 expression in CT26.WT and MC-38. The relative PD-L1 expression is the median fluorescence intensity of stained sample relative to secondary antibody alone. Data presented as mean ± S.E.M. of at least two independent experiments. Statistical significance calculated using student two-tailed t-test (*** p ≤ 0.001, * p ≤ 0.05).

## Discussion

The majority of cancer patients do not respond to ICT, and therefore there is a high need for novel rational strategies for overcoming resistance to this type of cancer immunotherapy. It has been suggested to look beyond checkpoint blockade, which may include combinatorial therapies that both are cytotoxic and eliminate cells that drive innate and adaptive immune suppression within the TME [43]. PD-L1 is highly expressed on tumor cells and immune cells in the TME, including DCs, macrophages, myeloid-derived suppressor cells (MDSCs), and Tregs [44]. All these immunosuppressive cell populations are, therefore, potential targets for cytotoxic attack with anti-PD-L1-aiming therapeutics.

This study demonstrates the *in vitro* proof-of-concept of a novel cytotoxic ICT strategy for the light-controlled targeting and elimination of PD-L1 positive cells. We show that PCI of an anti-human PD-L1 immunotoxin, anti-PD-L1-saporin, is a highly efficient strategy for eliminating cancer cells overexpressing PD-L1. In line with other studies, we confirm that PD-L1 is internalized in PD-L1+ cells [13–16]. Over time, the anti-PD-L1 mAb co-localizes with the lysosomotropic PCI photosensitizer fimaporfin/TPCS_2a_. In addition, co-localization of the PD-L1-tageting antibody with LysoTracker suggest that a fraction of internalized PD-L1 is transported to the lysosomes for degradation, and not only via recycling endosomes as previously shown [13]. After PCI, the fluorescence of TPCS_2a_ and AF488-labeled PD-L1 antibody becomes diffuse, indicating endo-/lysosomal escape and cytosolic release. Our results indicate that synergistic cytotoxic effects using PCI of the anti-PD-L1-targeting immunotoxin in the PD-L1 overexpressing human triple-negative breast cancer cell line MDA-MB-231 can be achieved in picomolar to nanomolar concentrations in MDA-MB-231 cells. Our data also provide evidence of specificity of the immunotoxin as treatment with unconjugated saporin did not induce any cytotoxic responses as compared to anti-PD-L1-saporin in the MDA-MB-231 cells. Moreover, PCI of anti-PD-L1-saporin was 50-fold more potent in reduction of cell viability than PCI of saporin. Lack of PCI-enhanced efficacy of anti-PD-L1-saporin was demonstrated in the PD-L1-negative control cell line MDA-MB-453, further validating the PD-L1-specificity of the immunotoxin.

Several groups have demonstrated the rationale of cytotoxic antibody-based targeting of immune checkpoints. One of the earliest reports by Tazzari *et al.* evaluated a single-chain fragment variable antibody (scFv) against CTLA-4 linked to saporin. The efficacy was demonstrated on activated T lymphocytes *in vitro* and *in vivo* as a strategy to prevent acute graft versus host disease (GVHD) and transplanted organ rejection [45]. Targeting of this CTLA-4-based immunotoxin was also suggested as a possible strategy for cancer treatment; however, preclinical data indicate that CTLA-4 has a significant role in regulating T cell responses to self-antigens [2], and systemic elimination of CTLA-4 positive cells may cause a high risk for severe adverse effects and even death. Moreover, clinical observations in cancer patients treated with anti-human CTLA-4 mAbs have shown that immune-related adverse events (irAEs) commonly occur [2,46].

The CTLA-4 or PD-1/PD-L1 axis regulates T-cell activation; however, these immune checkpoints modulate the T-cell immune response at different levels. CTLA-4 controls T cell activation in the lymphoid organs, whereas PD-1 is a negative regulator of T-cell primarily within the peripheral tissues and in the TME through interaction with PD-L1 [1,47]. As PD-L1 is mainly expressed on cancer cells or stromal immunosuppressive cells in the TME, targeting the PD-1/PD-L1 axis provides a more restricted spectrum of adverse effects, as demonstrated in clinical studies [47], and is, therefore, suitable for PCI-targeting. Moreover, with a few exceptions, normal human tissues rarely express PD-L1 protein on the cell surface [2]. Given that the PCI method enables spatiotemporal controlled drug delivery [27], we suggest that PCI-based cytotoxic targeting of PD-L1 positive cells in the TME of unresectable tumors has the potential of being both a safe and efficient method for targeting of immune suppressive cells due to; 1) The combo is non-toxic in dark and dependent on *in situ* light-controlled activation; 2) The PCI-photosensitizer accumulates more in malignant than in normal tissues [28,48]; 3) The anti-PD-L1 antibody provide specific targeting and uptake of the toxin cargo in PD-L1+ cells. However, this needs to be confirmed *in vivo*.

Li *et al.* provided evidence of an anti-human PD-L1 antibody conjugated to a potent anti-mitotic drug in syngeneic models with tumor cells expressing human PD-L1 [14]. The antibody-drug conjugate was shown to internalize; however, targeting the glycosylated domain of PD-L1 is crucial for intracellular uptake [14]. PD-L1 antibody conjugated to the chemotherapy drug doxorubicin and doxorubicin nanoparticles have also shown promising results [49,50].

Interestingly, the principle of a PD-L1 targeting immunotoxin, based on cucurmosin, has been demonstrated by Zhang *et al.* [51]. As saporin used in our study, cucurmosin is also a type I ribosome-inactivating protein (RIP) that potently inhibits protein synthesis but lacks an effective mechanism to enter cytosol upon endocytosis [52]. The FDA-approved anti-PD-L1 mAb durvalumab, commercially known as Imfinzi® by AstraZeneca, was conjugated to cucurmosin. The immunotoxin was highly potent in the picomolar range *in vitro*. The immunotoxin’s efficacy was further evaluated in an immunodeficient model. In the immunotoxin-treated animals, an initial tumor growth delay was observed. However, poor tumor penetration due to the size was identified as an obstacle. The size of the immunotoxin by Zhang *et al.* was approximately 230 kDa. Of relevance, the anti-PD-L1-saporin used in this study is 295 kDa. Based on our group’s previous experience, antibody-drug-conjugates of this size combined with PCI have indicated modest anti-tumor activity after PCI *in vivo*. The relatively large molecular size of these antibodies has been suggested to reduce the tumor penetration capacity and thereby therapeutic outcome [38,53]. One way of increasing tumor penetration capacity of immunotoxins is to replace the full-size receptor/ligand-targeting antibody with a scFv. We previously demonstrated that this strategy was, in combination with PCI, a specific and efficient anti-tumor strategy for targeting CSPG4-expressing melanoma and triple-negative breast cancer cells [37,54].

Moreover, in the context of *in vivo* preclinical testing, the use of anti-PD-L1-saporin in the current study is limited to human PD-L1 expressing cancer cells. Previous studies demonstrate that T cells are essential to achieve curative PCI effects [55,56]. We propose that PCI-based intracellular delivery of PD-L1 targeting immunotoxins is a feasible strategy to locally eradicate PD-L1 positive tumor cells and have the potential to eliminate immune-suppressive stromal cells expressing PD-L1 on their surface in the TME. We, therefore, aim to develop a murine version of the immunotoxin with a favorable size as this will be necessary to evaluate this strategy’s full potential in immunocompetent syngeneic tumor models.

In conclusion, we demonstrate for the first time that PCI promotes strong cytotoxic effects of a PD-L1 targeting immunotoxin in PD-L1 expressing cancer cells. PCI of the immunotoxin is superior to PCI of the non-targeted toxin or the immunotoxin alone, suggesting that this light-controlled, combinatorial cytotoxic ICT have the potential of being a specific, safe and potent therapeutic strategy to target and eradicate PD-L1 expressing immunosuppressive cells in the TME. Further preclinical validation of the principle is encouraged.

## Author Contributions

J.J.W.W. carried out the experiments, processed the experimental data, performed analysis, drafted the manuscript, and designed the figures. P.K.S conceived the original idea, provided funding and resources, supervised the project, and contributed to the final version of the manuscript.

## Acknowledgments

We would thank PCI Biotech for providing us with the PCI photosensitizer fimaporfin/TPCS_2a_ and Advanced Targeted Systems (ATS) for providing the anti-PD-L1-saporin immunotoxin. We would also like to thank the Hospital Pharmacy at Oslo University Hospital, Rikshospitalet, for providing us with unused atezolizumab (Tecentriq) from the Cancer Clinic.

